## Supplemental Figure 1 for "SARS-CoV-2 spike glycoprotein vaccine candidate NVX-CoV2373 elicits immunogenicity in baboons and protection in mice"

**Authors and Affiliations:**

Jing-Hui Tian[1 #], Nita Patel[1 #], Robert Haupt[2 #], Haixia Zhou[1], Stuart Weston[2], Holly Hammond[2], James Lague[2], Alyse D. Portnoff[1], James Norton[1], Mimi Guebre-Xabier[1], Bin Zhou[1], Kelsey Jacobson[1], Sonia Maciejewski[1], Rafia Khatoon[1], Malgorzata Wisniewska[1], Will Moffitt[1], Stefanie Kluepfel-Stahl[1], Betty Ekechukwu[1], James Papin[3], Sarathi Boddapati[4], C. Jason Wong[4], Pedro A. Piedra[5], Matthew B. Frieman[2], Michael J. Massare[1], Louis Fries[1], Karin Lövgren Bengtsson[6], Linda Stertman[6], Larry Ellingsworth[1], Gregory Glenn[1], and Gale Smith[1] *

[1]Novavax, Inc. 21 Firstfield Road, Gaithersburg, MD, 20878, USA. (J.H.T.), (N.P.), (H.Z.), (A.D.P.), (J.M), (M.G.X.), (B.Z.), (K.J.), (S.M.), (R.K.), (M.W.), (W.M.), (S.K.S.), (B.E.), (M.J.M.),

 (L.F.), (L.E.), (G.G.), (G.S.)

[2]University of Maryland, School of Medicine, 685 West Baltimore St, Baltimore, MD 21201, USA.. (M.B.F., R.H., S.W., H.H.)

[3]University of Oklahoma, Health Sciences Center, Department of Pathology, Division of Comparative Medicine, 940 Stanton L. Young, BMS 203, Oklahoma City, OK, 73104 USA. (J.P.)

[4]Catalent Paragon Gene Therapy, 801 West Baltimore Street, Baltimore, MD 21201. USA. (S.B.), (C.J.W.)

[5]Department of Molecular Virology and Microbiology, and Pediatrics, Baylor College of Medicine, Houston, Texas. (P.A.P.)

[6]Novavax AB, Kungsgatan 109, Uppsala, SE-753 18, SE. (K.L.B.), (L.S.)

[#]JHT, NP, RH and each contributed equally as co-lead authors.

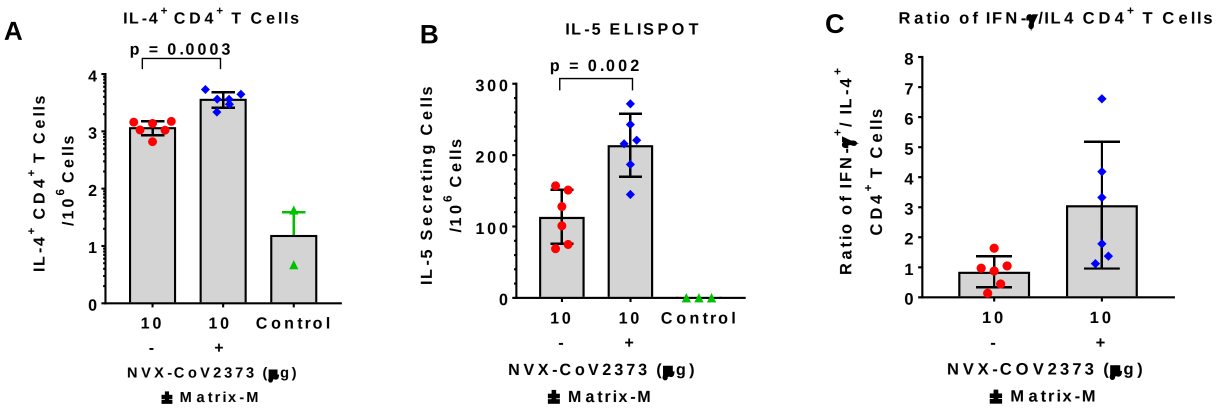


**Supplementary Figure 1. Intracellular staining (ICCS) and ELISPOT detection of type 2 cytokines in immunized mice.** Groups of mice were immunized with NVX-CoV2373 with and without 5 µg Matrix-M adjuvant with 2-doses spaced 21-days apart and splenocytes collected 7-days after the second immunization and stimulated with NVX-CoV2373 protein. (**A**) ICCS of IL-4$^+$ CD4$^+$ T cells. (**B**) IL-5-secreting splenocytes determined by ELSPOT analysis. Data show the mean ± SD. (**C**) Ratio of antigen-specific IFN-γ$^+$ to IL-4$^+$ CD4$^+$ T cells in spleens of mice immunized with NVX-CoV2373 with and without Matrix-M adjuvant.
